# Antioxidant properties of dihydroxy B-ring flavonoids modulate circadian amplitude in *Arabidopsis*

**DOI:** 10.1101/2025.03.09.641856

**Authors:** Evan S. Littleton, Sherry B. Hildreth, Shihoko Kojima, Brenda S.J. Winkel

## Abstract

Flavonoids are an abundant specialized metabolite produced by plants for a range of functions, including pigmentation, hormonal signaling, UV protection, and drought tolerance. We previously showed that flavonoids also influence the circadian clock in *Arabidopsis*. Here, we report that the antioxidant properties of dihydroxy B-ring flavonoids is responsible for regulating the amplitude of the core clock gene luciferase reporter, *TOC1:LUC.* We found the amplitude of *TOC1:LUC* rhythms correlates with the cellular H_2_O_2_ content in flavonoid-deficient seedlings. Moreover, reducing production of reactive oxygen species rescued the elevated *TOC1:LUC* amplitude in flavonoid-deficient seedlings, whereas reducing auxin transport rate, a known function of flavonoids, had no impact on *TOC1:LUC* amplitude. Interestingly, Ca^2+^ levels in the chloroplast, but not the cytosol, were also altered in flavonoid-deficient seedlings, hinting at retrograde signaling as a possible mechanism of flavonoid-mediated changes in clock amplitude. This study advances our understanding of the relationship between flavonoids and the circadian clock in plants and deepens our understanding of the mechanisms underlying this interaction.

## Introduction

The pervasiveness of circadian rhythms across the tree of life is indicative of the evolutionary benefit provided by the ability to predict and prepare for a changing daily environment (Jabbur & Johnson, 2022; Laosuntisuk *et al*, 2023; Oravec & Greenham, 2022). In plants, circadian rhythms control nearly all aspects of physiology, including cellular metabolism, photosynthetic activity, growth, stomatal opening, defense from pathogens and herbivores, leaf movement, and flowering (Creux & Harmer, 2019; Karapetyan & Dong, 2018; Venkat & Muneer, 2022). Plants that do not match the pace of their internal clock to the external environment, either through altered circadian period or arrhythmia, have decreased chlorophyll content, CO_2_ assimilation, and biomass (Dodd *et al*, 2005). In addition to the period/pace of the clock, there is now growing evidence for the importance of amplitude/robustness, as a robust clock has been co-selected with agriculturally-important crop traits, particularly flowering time. The recently-named field of “chronoculture” proposes to target the plant circadian clock through genetic modification or timing-based agricultural practices to further enhance crop productivity and agricultural sustainability (Creux & Harmer, 2019; Steed *et al*, 2021).

At the cellular level, circadian rhythms are generated by a transcription-translation feedback loop (TTFL) consisting of multiple complex interactions between transcription factors (Creux & Harmer, 2019; Laosuntisuk *et al*., 2023). In plants, the evening gene, *TOC1*, and the morning genes, *CCA1/LHY*, are central players in a network of overlapping feedback loops that generates a robust 24-hour cycle of transcription and translation regulating thousands of downstream genes (Harmer *et al*, 2000; Liebelt *et al*, 2019; Nagel *et al*, 2015). While the TTFL is predominantly synchronized by external input from the environment such as light, temperature, and humidity (Millar, 2004; Mwimba *et al*, 2018; Salomé & McClung, 2005), recent studies have shown the importance of inputs from internal sources as well; clock gene amplitude, period, or phase can be modified through photosynthetic sugars (Dalchau *et al*, 2011; Frank *et al*, 2018; Haydon *et al*, 2017; Haydon *et al*, 2013; Román *et al*, 2021), cytosolic free calcium (Martí Ruiz *et al*, 2018), and reactive oxygen species (ROS) (Román *et al*., 2021; Zhou *et al*, 2015). These studies showcase the innate sensitivity and adaptability of the plant clock to both environmental cues and internal cellular changes, as well as highlighting the importance of reciprocal signaling from metabolites as a critical aspect of how the TTFL is regulated.

We recently showed that a loss of flavonoid biosynthesis in *Arabidopsis* modulates clock gene expression, affecting both transcript levels and the amplitude of *TOC1:LUC* rhythms, without significantly altering phase or period (Hildreth *et al*, 2022). This influence on clock function adds to a long list of well-established roles for flavonoids in plants, including functions in pigmentation, development, and stress protection. Many of these roles have been attributed to the strong antioxidant potential of flavonoids, particularly the dihydroxy B-ring forms such as quercetin and cyanidin (Fig. EV1; Agati *et al*, 2012; Agati *et al*, 2020; Daryanavard *et al*, 2023; Gayomba & Muday, 2020; Hernández *et al*, 2009; Nakabayashi *et al*, 2014; Xu *et al*, 2017). A recent study has uncovered a role for superoxide as a metabolic signal that affects the amplitude of clock gene expression (Román *et al*., 2021), prompting us to investigate the possibility that a loss of antioxidant activity is responsible for the elevated *TOC1:LUC* amplitude in flavonoid mutants. Here, we show that a deficiency of dihydroxy B-ring flavonoids in *Arabidopsis* seedlings enhances the amplitude of *TOC1:LUC,* predominantly via elevated ROS level. Using biochemical and genetic approaches we begin to elucidate the mechanisms underlying the ROS-dependent modulation of clock gene expression by flavonoids.

## Results

### Exogenously-supplied flavonoids affect *TOC1* promoter activity

In a previous study, we found that bioluminescence output from *TOC1:LUC* had a higher amplitude in *tt4-11* seedlings relative to wild-type controls (Hildreth *et al*, 2020). The *tt4-11* line lacks all flavonoids due to a T-DNA insertion in the gene encoding chalcone synthase (CHS), the first enzyme in the flavonoid biosynthetic pathway (Bowerman *et al*, 2012; Fig. EV1). Bioluminescence output from *TOC1:LUC* also had a higher amplitude in *tt7-5* mutants, which lack the enzyme, flavonoid 3’ hydroxylase (F3’H), also due to a T-DNA insertion (Bowerman *et al*., 2012) and can therefore produce monohydroxy but not dihydroxy B-ring flavonoids. Furthermore, we showed that supplementing the medium with the dihydroxylated flavonol, quercetin, affected the amplitude of the *CCA1:LUC* reporter in both wild type and *tt4-11* seedlings. These data led us to hypothesize that dihydroxy B-ring flavonoids play a predominant role in modulating circadian amplitude.

To test this hypothesis, we first focused on the impact of exogenously applied flavonoids on expression of the *TOC1:LUC*, as this reporter exhibits enhanced amplitude in flavonoid mutant lines, *tt4-11,* and *tt7-5* (Hildreth *et al*., 2022). Seeds for the Col-0 wild-type, *tt4-11,* and *tt7-5* lines containing the *TOC1:LUC* reporter were sown on medium containing naringenin, the first intermediate in flavonoid biosynthesis (Fig. EV1). After six days incubation in a 12 h light:12 h dark (LD) cycle, the seedlings were transferred to constant darkness (DD) for luminescence measurements. For both Col-0 and *tt4-11,* the amplitude of *TOC1:LUC* bioluminescence was slightly enhanced in the presence of 1 µM naringenin compared to the control (DMSO), while 10 µM naringenin had no effect (Figs. 1A-B and EV2A; File S1). At 100 µM, however, *TOC1:LUC* amplitude decreased in Col-0 and decreased even more drastically in *tt4-11.* No changes were observed in the period of *TOC1:LUC* luminescence with these treatments (Fig. EV3A), consistent with our previous report (Hildreth *et al*., 2022). In contrast, treatment with 100 µM naringenin did not have any effect on the *TOC1:LUC* amplitude in *tt7-5*, which cannot convert naringenin to dihydroxy B-ring flavonoids (Fig. 1C-D). These data support our hypothesis that the dihydroxy B-ring forms of flavonoids are primarily responsible for the changes in the amplitude of *TOC1:LUC* rhythms.

**Figure 1:**
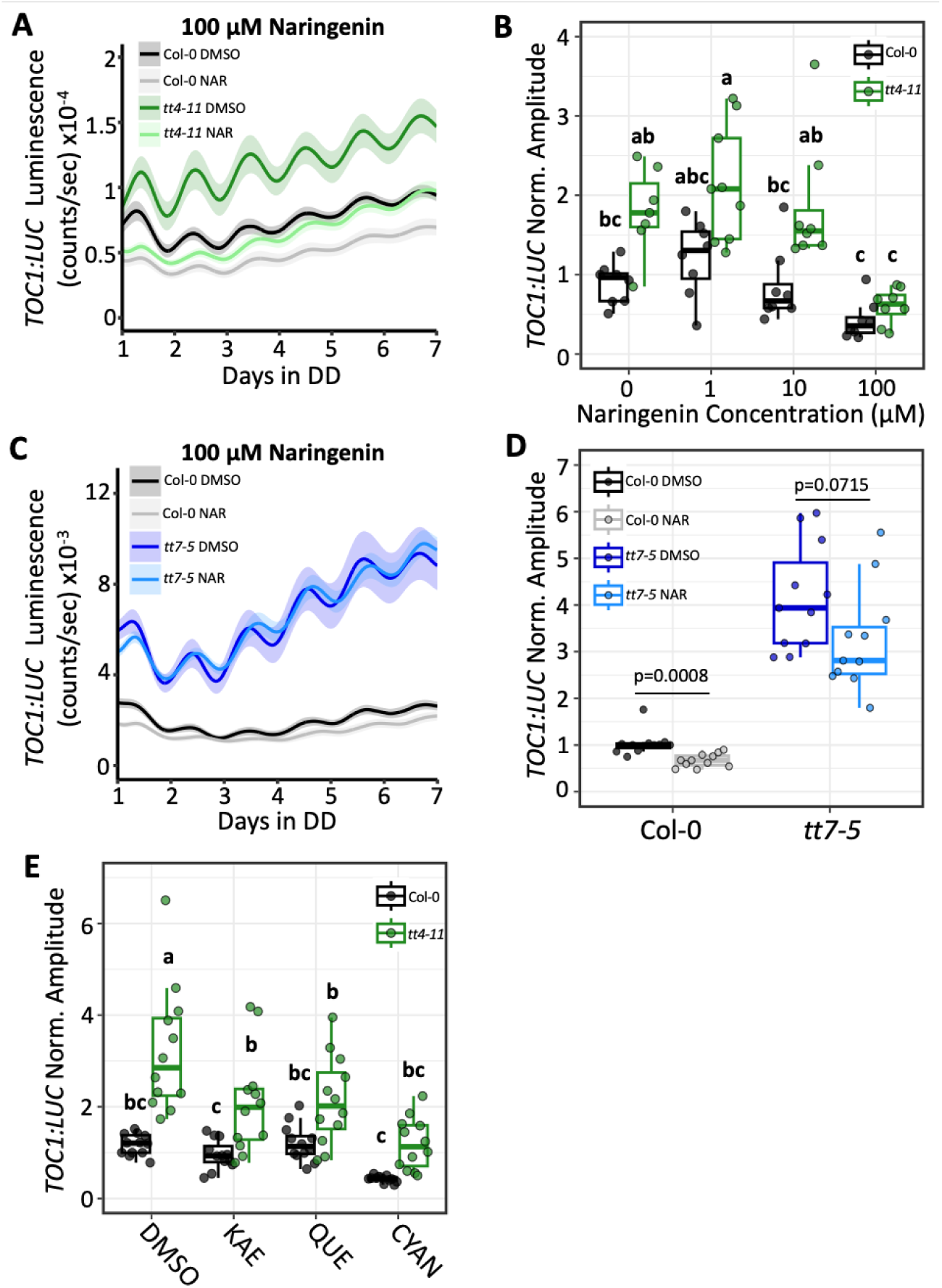
Effects of exogenous flavonoids on *TOC1:LUC* amplitude in wild-type and flavonoid-deficient lines. (A) Bioluminescent output of *TOC1:LUC* in Col-0 or *tt4-11* seedlings grown on 0.1% DMSO (black: Col-0, green: *tt4-11*) or 100 µM naringenin (NAR) (grey: Col-0, light green: *tt4-11*) (n=8 biological replicates from two independent experiments). (B) Amplitude of *TOC1:LUC* (black: Col-0, green: *tt4-11*) with 0, 1, 10, or 100 µM NAR. Values normalized to Col-0 0 µM, which was set to 1. (C) Bioluminescent output of *TOC1:LUC* in Col-0 or *tt7-5* seedlings grown on 0.1% DMSO (black: Col-0, blue: *tt7-5*) or 100 µM NAR (grey: Col-0, light blue: *tt7-5*) (n=10-11 biological replicates from three independent experiments). (D) Amplitude of *TOC1:LUC* in Col-0 or *tt7-5* grown on 0.1% DMSO or 100 µM NAR. Values normalized to Col-0 DMSO, which was set to 1. (E) Amplitude of *TOC1:LUC* (black: Col-0, green: *tt4-11*) with 0.1% DMSO or 100 µM kaempferol (KAE), quercetin (QUE), or cyanidin (CYAN) (n=12 biological replicates from three independent experiments). Values normalized to Col-0 DMSO, which was set to 1. P-values in D are calculated from two-tailed Student’s T-test (Col-0; unequal variance, *tt7-5*; equal variance). Letters in B and E represent grouping from one-way ANOVA followed by Tukey post-hoc test with significance cutoff of p<0.05. In B, D, and E, boxplot midlines represent median value (Q2), lower and upper line represent 25^th^ (Q1) and 75^th^ percentiles (Q3), respectively. Whiskers represent range of data within 1.5 interquartile range (Q3-Q1) from Q1 or Q3. Solid line and shading in A and C represent mean ± SEM.

We next tested the effects of the predominant flavonoid types produced in *Arabidopsis* on *TOC1:LUC* expression. We found that at 100 µM kaempferol and quercetin (monohydroxy and dihydroxy B-ring flavonols, respectively) both lowered *TOC1:LUC* amplitude in *tt4-11,* while cyanidin (a dihydroxy B-ring anthocyanidin), fully reduced amplitude to wild-type control levels (Figs. 1E and EV2B). In contrast to our observations for *tt4-11,* exogenous flavonoids had only a minor effect on *TOC1:LUC* amplitude in Col-0, although cyanidin did appear to consistently reduce reporter amplitude relative to wild-type controls. Although the similar effects of the two flavonols was unexpected, this may be the result of conversion of kaempferol to quercetin through the action of F3’H (e.g., Ueyama *et al*, 2002; Zhou *et al*, 2016), which remains fully functional in *tt4-11.* It also appears that cyanidin has an even greater impact on *TOC1:LUC* amplitude in *tt4-11* than either flavonol, although differential uptake or conversion/modification cannot be ruled out at this stage. Consistent with the effects observed with naringenin, the period of *TOC1:LUC* expression remained unchanged under all conditions (Fig. EV3B). Overall, these findings show that exogenous application of different flavonoid subtypes, and particularly cyanidin, a dihydroxy B-ring anthocyanidin, can restore the enhanced amplitude in flavonoid-deficient seedlings back to wild-type levels.

### Few flavonol glycosides exhibit rhythmicity across the day-night cycle

The expression of flavonoid biosynthetic genes is highly rhythmic at the mRNA level (Harmer *et al*., 2000; Liebelt *et al*., 2019; Nagel *et al*., 2015), however, little is known about the accumulation patterns of the pathway’s end products across the day-night cycle in seedlings. Since there is often a poor correlation between the transcriptome and metabolome even within the same pathway (Hildreth *et al*., 2020), we asked whether specific flavonoids exhibit rhythmic patterns that could underlie modulation of clock gene amplitude. To this end, we used untargeted LC-MS/MS to examine metabolite profiles over 24h in 7-day-old Col-0 seedlings grown under a 12h:12h LD cycle, where zeitgeber time 0 and 12 represent the onset and offset of light, respectively. Of the eight flavonoid glycosides identified in our dataset (predominantly kaempferol, quercetin, and isorhamnetin glycosides), only the monohydroxylated flavonol, kaempferol deoxyhexose, and the dihydroxylated flavonol, isorhamnetin hexose deoxyhexose, showed a rhythmic pattern of accumulation at a statistically-significant level. All other flavonoids showed a weak diurnal pattern, but with very low amplitude and statistically-insignificant rhythmicity (Fig. 2; File 2). These observations in seedlings are consistent with a recent finding that flavonoid glycosides do not accumulate rhythmically in LD or LL in *Arabidopsis* leaves (Rivière *et al*, 2024). However, it remains possible that specific flavonoids responsible for modulating clock behavior have rhythmic accumulation patterns that are only discernable at the cellular or organellar level.

**Figure 2:**
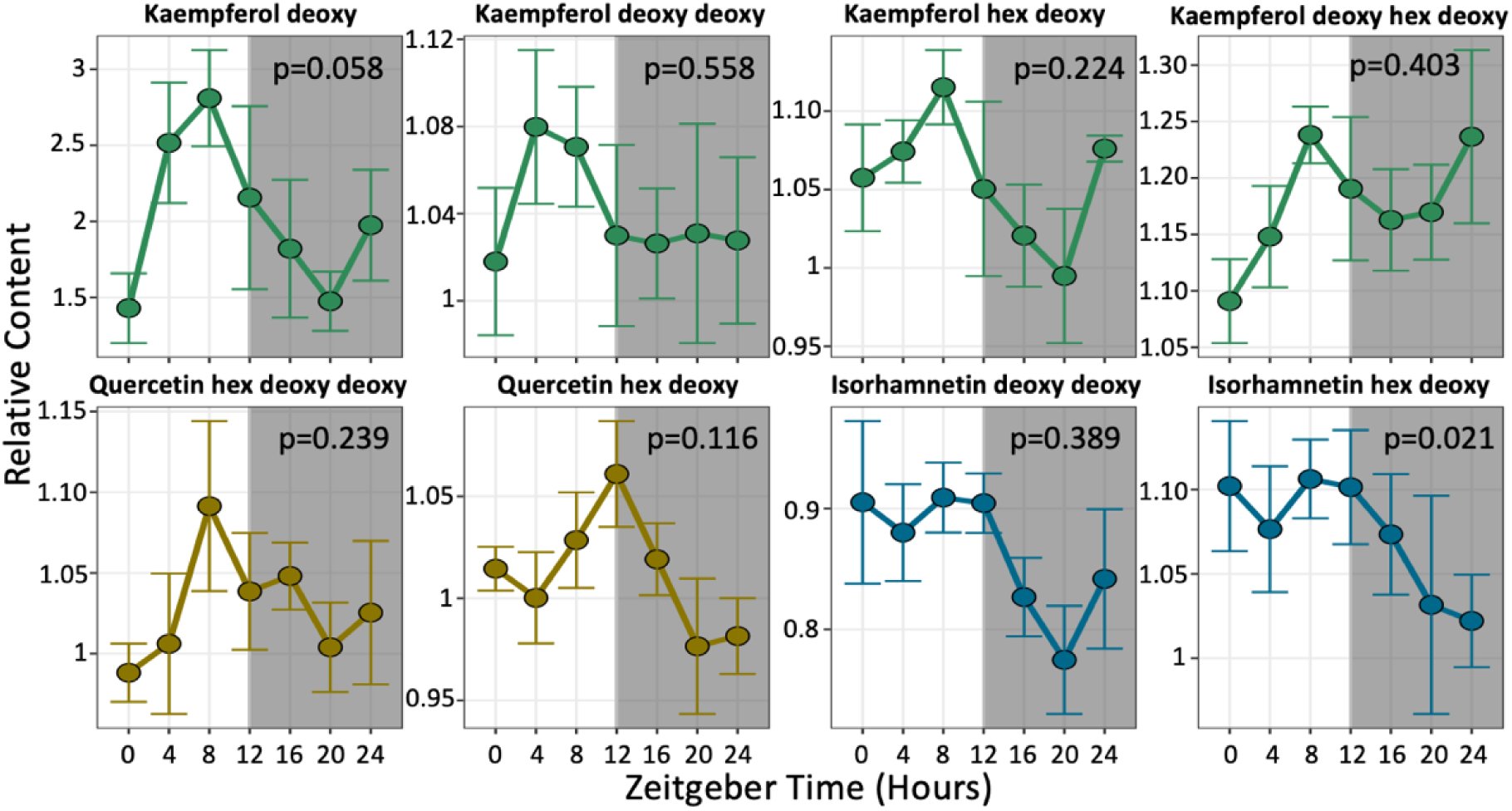
Flavonoid accumulation patterns in wild-type seedlings. Methanolic extracts from 7-day-old Col-0 plants grown under LD were examined by LC-MS (n=4-5 biological replicates for each time point). All the data represent mean ± SEM. Values at zeitgeber time 0 (onset of light) were set as 1. P-values for rhythmicity are calculated from MetaCycle with algorithms JTK, LS, and ARS.

### Effects of flavonoids on *TOC1:LUC* amplitude do not involve auxin signaling

A well-established function of flavonoids is the inhibition of auxin transport (Brown *et al*, 2001; Buer & Muday, 2004; Lewis *et al*, 2011). This is believed to occur through interactions of flavonols such as quercetin and kaempferol with PIN-FORMED (PIN) or ABCB transporters responsible for intracellular movement of auxin (Daryanavard *et al*., 2023; Teale *et al*, 2021). Interestingly, previous studies found that several clock gene reporters, including *TOC1:LUC*, exhibited reduced amplitude when seedlings were grown in the presence of indole-3-acetic acid (IAA), the most common form of auxin found in plants (Covington & Harmer, 2007; Hanano *et al*, 2006). To test the possibility that the increased auxin transport, and resulting altered distribution, in *tt4-11* (Buer & Muday, 2004) causes the elevation in *TOC1:LUC* amplitude, we grew seedlings in the presence of N-1-naphthylphthalamic acid (NPA), which inhibits auxin transport by interacting with PINs and ABCB transporters (Teale *et al*., 2021). When we used a range of concentrations that were previously shown to reduce root gravitropism and growth (Brown *et al*., 2001; Rashotte *et al*, 2000), NPA had no significant impact on *TOC1:LUC* amplitude or period in either Col-0 or *tt4-11* seedlings (Figs. 3A-B, EV2C, and EV3C; File S1). Together, these findings indicate that auxin does not play a role in mediating the effects of flavonoids on the amplitude of *TOC1:LUC*.

**Figure 3:**
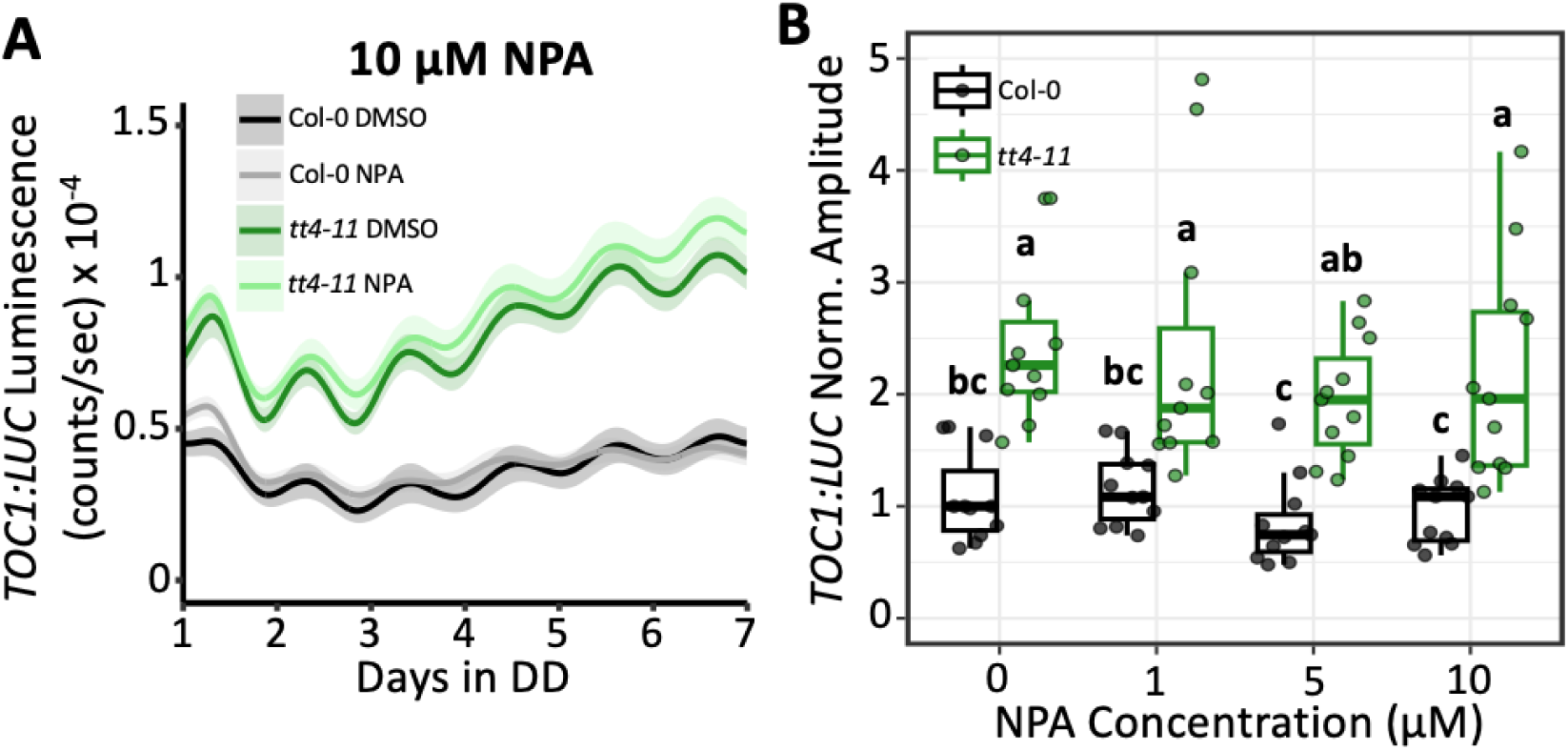
Effects of auxin transport inhibitor on *TOC1:LUC* amplitude. (A) Bioluminescent output of *TOC1:LUC* in Col-0 or *tt4-11* seedlings grown on 0.1% DMSO (black: Col-0, green: *tt4-11*) or 10 µM NPA (grey: Col-0, light green: *tt4-11*). Solid line and shading represent mean ± SEM (n=11 biological replicates from three independent experiments). (B) Amplitude of *TOC1:LUC* (black: Col-0, green: *tt4-11*) grown on 0, 1, 5, or 10 µM NPA. Values normalized to Col-0 0 µM, which was set to 1. Letters in B represent grouping from one-way ANOVA followed by Tukey post-hoc test with significance cutoff of p<0.05. Boxplot midlines represent median value (Q2), lower and upper line represent 25^th^ (Q1) and 75^th^ percentiles (Q3), respectively. Whiskers represent range of data within 1.5 interquartile range (Q3-Q1) from Q1 or Q3.

### *TOC1:LUC* amplitude correlates with H_2_O_2_ content

Due to the strong antioxidant capabilities of dihydroxy B-ring flavonoids (Agati *et al*., 2012; Agati *et al*., 2020; Nakabayashi *et al*., 2014) and ability of ROS to modulate circadian amplitude (Román *et al*., 2021; Zhou *et al*., 2015), we next asked whether the elevated amplitude of *TOC1:LUC* observed in *tt4-11* is driven by elevated ROS levels. To test this, we first measured H_2_O_2_ levels in *tt4-11* and Col-0 seedling lysates across 24-hour cycles under different light conditions (Fig. 4A). Consistent with previous reports of elevated H_2_O_2_ and O_2_^-^ in flavonoid-mutant seedlings (Chapman & Muday, 2021; Xu *et al*., 2017), we found that in LD (12h light/12h dark), the H_2_O_2_ levels were approximately 3-fold higher in *tt4-11* than in wild-type seedlings at all time points (Fig. 4B). The levels of H_2_O_2_ also remained two- to three-fold higher in *tt4-11* following transfer from LD to constant light (LL) or constant darkness (DD) (Fig. 4C-D).

**Figure 4:**
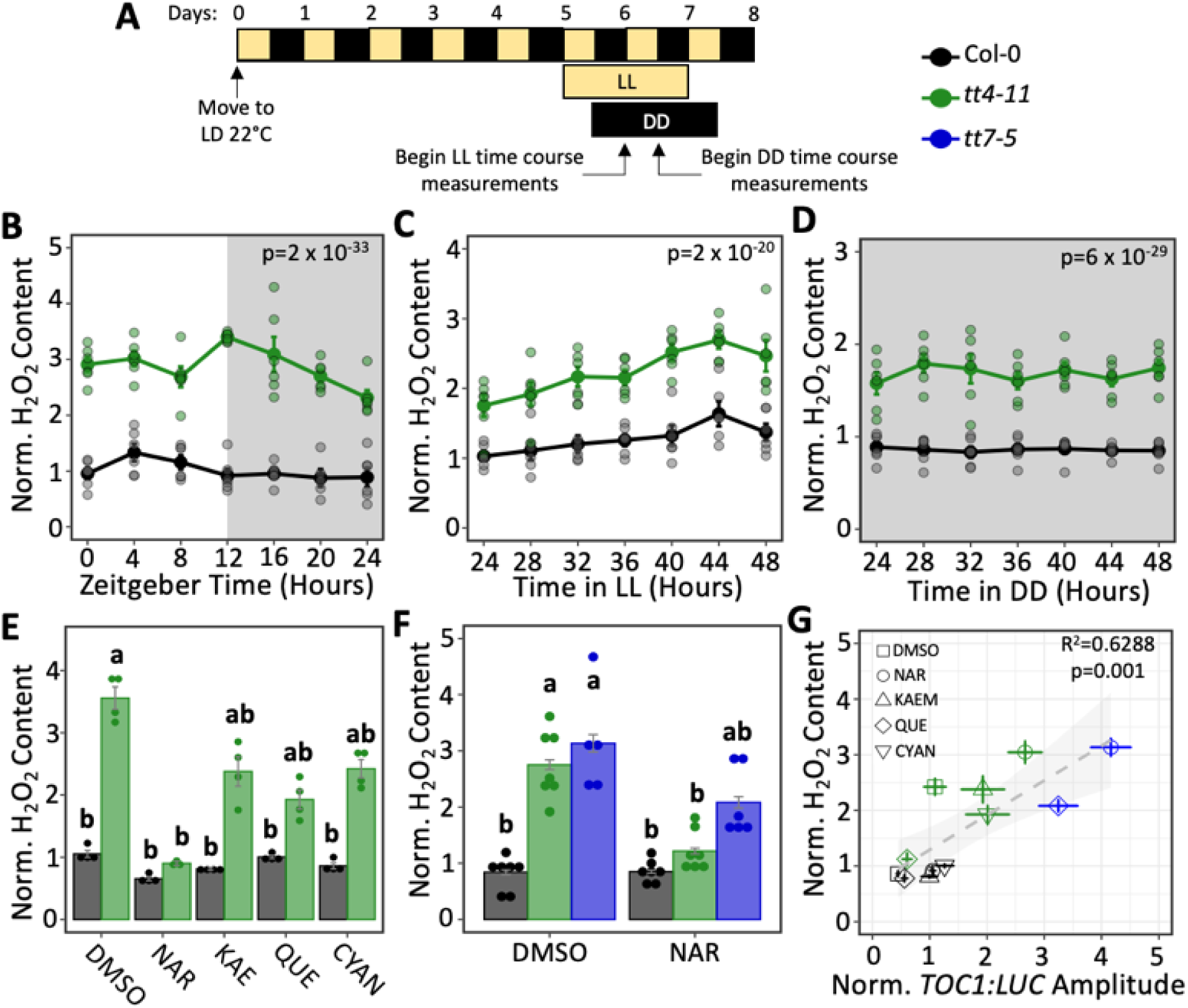
Correlation between H2O2 levels and *TOC1:LUC* amplitude. (A) diagram of experimental timeline for sample preparation and collection. (B) H2O2 content of 6-day old seedlings (black: Col-0, green: *tt4-11*) collected every 4 h for 24 h in LD (n=5-6 biological replicates/time point from two independent experiments). Values normalized to Col-0 zeitgeber time 0, which was set to 1 (C) H2O2 content of 6-day-old seedlings (black: Col-0, green: *tt4-11*) transferred to LL for 24 h and collected every 4 h for 24 h (n=6 biological replicates/time point from two independent experiments). Values normalized to Col-0 timepoint 24, which was set to 1 (D) H2O2 content of 6-day old seedlings (black: Col-0, green: *tt4-11*) transferred to DD for 24 h and collected every 4 h for 24 h (n=6 biological replicates/time point from two independent experiments). Values normalized to Col-0 timepoint 24, which was set to 1 (E) H2O2 content of 6-day-old seedlings (black: Col-0, green: *tt4-11*) grown on media containing DMSO or various flavonoids transferred to DD for 24 h before collection (n=3-4 biological replicates from two independent experiments). Values normalized to Col-0 DMSO, which was set to 1. (F) H2O2 content of 6-day old seedlings (black: Col-0, green: *tt4-11* blue: *tt7-5*) grown on media containing DMSO or NAR transferred to DD for 24 h before collection (n=5-7 biological replicates/time point from three independent experiments). Values normalized to Col-0 DMSO, which was set to 1. (G) Linear regression model between *TOC1:LUC* amplitude and H2O2 content. P-values in B-D calculated from two-way ANOVA. Letters in E-F represent grouping from one-way ANOVA followed by Tukey post-hoc test with significance cutoff of p<0.05. All data represent mean ± SEM.

We then asked whether exogenous flavonoids, which had an effect on the *TOC1:LUC* amplitude (Fig. 1), also influence H_2_O_2_ levels. The level of H_2_O_2_ in the wild-type seedlings was unchanged in the presence of NAR, KAE, QUE, or CYAN (Fig. 4E). In contrast, the level of H_2_O_2_ in *tt4-11* seedlings was slightly, but insignificantly, reduced by KAE, QUE, or CYAN, while it was fully restored to wild-type levels by NAR. To determine whether the dihydroxy B-ring forms of flavonoids are primarily responsible for modulating H_2_O_2_ levels, we examined the effects of exogenous flavonoids on H_2_O_2_ levels in *tt7-*5. In contrast to the effects in *tt4-11*, NAR only partially reduced the H_2_O_2_ level in the *tt7-5* seedlings relative to wild type (Fig. 4F). These results indicate that a lack of dihydroxy B-ring flavonoids is primarily responsible for the elevated H_2_O_2_ levels in flavonoid mutant lines.

To better determine whether the antioxidant activity of flavonoids is responsible for suppressing the amplitude of *TOC1:LUC*, we next performed a correlation analysis between *TOC1:LUC* amplitude and H_2_O_2_ levels across all three genotypes and four flavonoid treatment groups. Using a linear regression model, we saw a positive linear correlation between *TOC1:LUC* amplitude and H_2_O_2_ content (Fig. 4G), strongly suggesting that flavonoids, and particularly the dihydroxy B-ring forms, regulate the amplitude of *TOC1:LUC* via their antioxidant activities.

We then asked whether reducing ROS levels in *tt4-11* would lower the elevated *TOC1:LUC* amplitude in these seedlings. To this end we first used a chemical approach, treating seedlings with diphenyleneiodonium (DPI). This inhibitor of the NADPH-oxidases that generate O_2_^-^, a precursor of H_2_O_2_, at the plasma membrane, has previously been shown to strongly attenuate the enhancement of *TOC1:LUC* amplitude in wild-type seedlings in response to sucrose (Román *et al*., 2021). DPI treatment of *tt4-11* led to a significant dose-dependent reduction in *TOC1:LUC* amplitude, with 10 µM and 30 µM treatments resulting in amplitudes that were equal to or lower than, respectively, those in untreated wild-type seedlings (Figs. 5A-B and EV2D, File S1). DPI also decreased *TOC1:LUC* amplitude in Col-0, although the difference was not statistically significant. Similar to treatment with flavonoids, no changes in the period of *TOC1:LUC* expression were observed at any concentration of DPI (Fig. EV3D).

**Figure 5:**
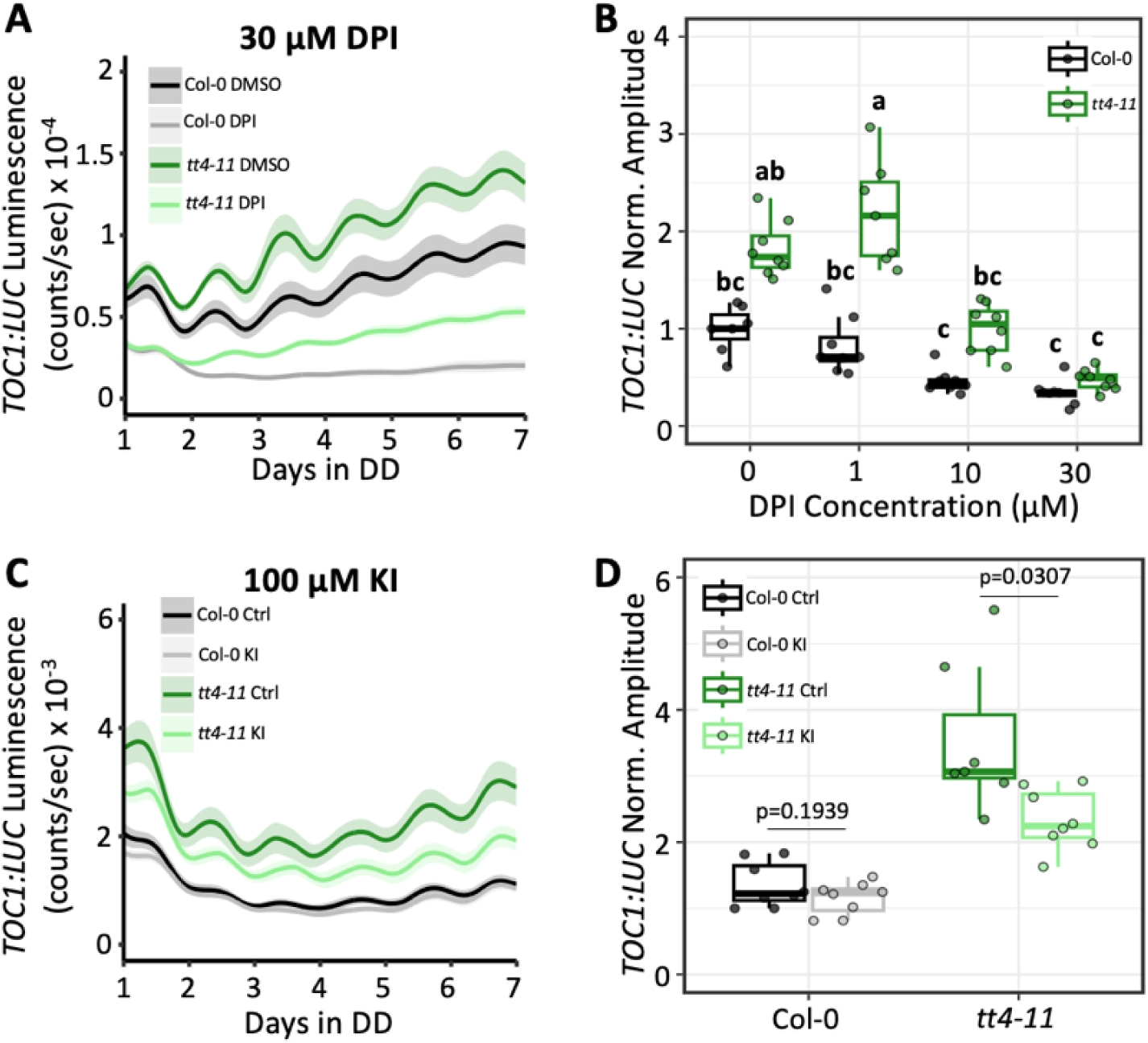
Effect of chemically-induced ROS reduction on *TOC1:LUC* amplitude in *tt4-11.* (A) Bioluminescent output of *TOC1:LUC* in Col-0 or *tt4-11* seedlings treated with 0.1% DMSO (black: Col-0, green: *tt4-11*) or 30 µM DPI (grey: Col-0, light green: *tt4-11*) (n=7-8 biological replicates from two independent experiments). (B) Amplitude of *TOC1:LUC* in seedlings (black: Col-0, green: *tt4-11*) treated with 0, 1, 10, or 30 µM DPI. Values normalized to Col-0 at 0 µM, which was set to 1. (C) Bioluminescent output of *TOC1:LUC* in seedlings (black: Col-0, green: *tt4-11*) grown on untreated media or 100 µM KI (grey: Col-0, light green: *tt4-11*) (n=7-8 biological replicates from two independent experiments). (D) Amplitude of *TOC1:LUC* in seedlings (black: Col-0, green: *tt4-11*) treated with 0 (Ctrl) or 100 µM KI. Values normalized to Col-0 Ctrl, which was set to 1. P-values in D calculated with two-tailed Student’s T-test with equal variance. Letters in B represent grouping from one-way ANOVA followed by Tukey post-hoc test with significance cutoff of p<0.05. In B and D, boxplot midlines represent median value (Q2), lower and upper line represent 25^th^ (Q1) and 75^th^ percentiles (Q3), respectively. Whiskers represent range of data within 1.5 interquartile range (Q3-Q1) from Q1 or Q3. Solid line and shading in A and C represent mean ± SEM.

We next tested another approach to reducing intracellular ROS, and treated the seedlings with the H_2_O_2_ scavenger, potassium iodide (KI) (Agati *et al*., 2020; Gayomba & Muday, 2020). At 100 µM KI, the amplitude of *TOC1:LUC* was reduced in *tt4-11* seedlings relative to untreated controls, whereas there was no impact in Col-0 (Fig. 5C-D). Together with the effects observed for treatment with DPI, these results show that the enhanced amplitude of *TOC1:LUC* in *tt4-11* can be lowered by reducing ROS levels either by inhibiting O_2_ ^-^ generation or by supplementing with H_2_ O_2_ scavengers. These findings provide further evidence that flavonoids suppress *TOC1:LUC* amplitude through their antioxidant properties.

### Modulation of *TOC1:LUC* amplitude by flavonoids does not involve the NPR1 receptor

We next attempted to gain mechanistic insights into how the antioxidant activity of flavonoids regulate the amplitude of *TOC1:LUC*. We first focused on NPR1 (non-expressor of pathogenesis-related gene 1), a master regulator of the plant immune response, and hypothesized that flavonoid modulation of H_2_O_2_ levels regulate the core clock through NPR1. This is because NPR1 controls the expression of *TOC1* and other clock genes via the plant’s redox state (Zhou *et al*., 2015), and because naringenin induces nuclear localization of NPR1 in a ROS-dependent manner (An *et al*, 2021). If our hypothesis was correct, we predicted that the changes in the amplitude of the *TOC1:LUC* reporter by exogenous flavonoids (Fig. 1) would be lost in the absence of NPR1. The amplitude of *TOC1:LUC* was lower in *npr1-3,* a NPR1 null line, than Col-0 seedlings, consistent with the previous report (Zhou *et al*., 2015). However, the amplitude and the basal level of *TOC1:LUC* was reduced in the presence of 100 µM naringenin relative to untreated controls (Fig. EV 4A-B), in both Col-0 and *npr1-3.* These data indicate that the effect of flavonoids on the amplitude of *TOC1:LUC* is independent of NPR1.

### Flavonoids may influence clock gene expression through Ca^2+^ signaling in chloroplasts

We next focused on cytosolic free calcium level ([Ca^2+^]_cyt_) and hypothesized the increased H_2_O_2_ content in *tt4-11* (Fig. 4) alters daily [Ca^2+^]_cyt_ rhythms, which in turn lead to the elevated amplitude of *TOC1:LUC*. This is because changes to [Ca^2+^]_cyt_ alter the expression of clock genes through a pathway involving TOC1 (Martí Ruiz *et al*., 2018). Moreover, ROS increases [Ca^2+^]_cyt_ in plants by altering the activity of Ca^2+^ channels/transporters (Fichman *et al*, 2022; Mazars *et al*, 2010; Mori & Schroeder, 2004; Ravi *et al*, 2023). To address this, we used the *Arabidopsis* MAQ2 line expressing cytosolic aequorin, a luminescent Ca^2+^ biosensor (Johnson *et al*, 1995; Knight *et al*, 1991), and crossed this with Col plants carrying the *tt4-2* allele (Bennett *et al*, 2006) to circumvent silencing effects from the T-DNA insertion in *tt4-11* (Daxinger *et al*, 2008). *tt4-2* also lacks a functional CHS enzyme, in this case due to a point mutation that disrupts pre-mRNA splicing (Burbulis *et al*, 1996). When we monitored the bioluminescent output from cytosolic aequorin (MAQ2) using six-day-old LD grown seedlings transferred to DD, we saw no changes in the amplitude of circadian [Ca^2+^]_cyt_ oscillations in *tt4-2* compared to Col-0 (Fig. 6A-B).

**Figure 6:**
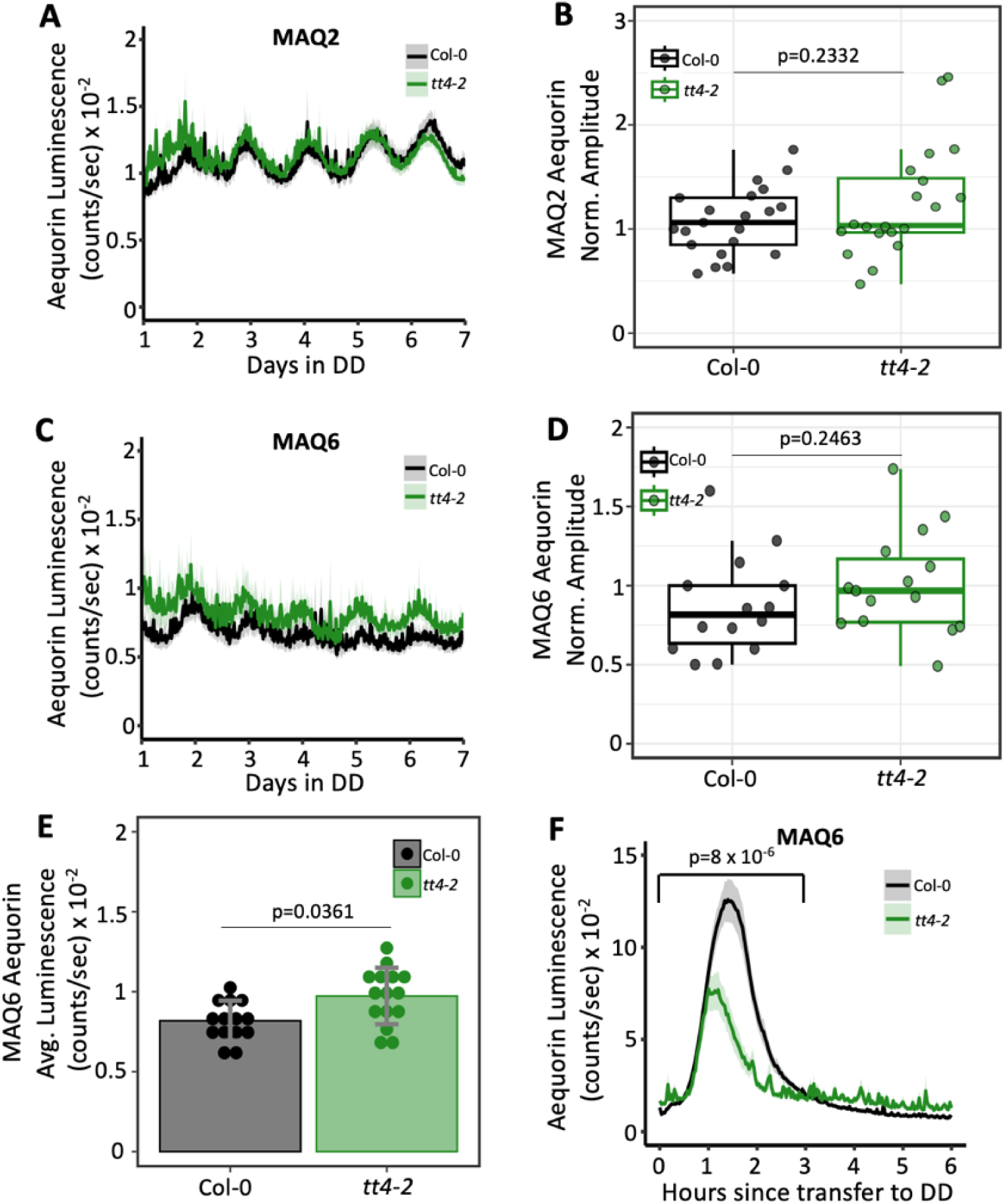
Flavonoid deficiency alters Ca^2+^ levels in the chloroplast. (A) Bioluminescence output of cytosolic aequorin (MAQ2) in Col-0 (black) or *tt4-2* (green) (n=20-21 biological replicates from two independent experiments). (B) Amplitude of cytosolic aequorin rhythms in Col-0 or *tt4-2* normalized to Col-0, which was set to 1. P-value calculated from two-tailed Student’s T-test with unequal variance. (C) Bioluminescence output of chloroplast targeted (MAQ6) aequorin in Col-0 (black) or *tt4-2* (green) (n=14-15 biological replicates from two independent experiments). (D) Amplitude of chloroplast-targeted aequorin rhythms in Col-0 or *tt4-2* normalized to Col-0, which was set to 1. P-value calculated from two-tailed Student’s T-test with equal variance. (E) Average luminescence of chloroplast-targeted aequorin from days 1-7 in DD. P-value calculated from two-tailed Student’s T-test with unequal variance. (F) Bioluminescence output of chloroplast targeted aequorin for 6 hours post-transfer from LD to DD. P-value calculated from two-way ANOVA between hours 0-3. In B and D, boxplot midlines represent median value (Q2), lower and upper line represent 25^th^ (Q1) and 75^th^ percentiles (Q3), respectively. Whiskers represent range of data within 1.5 interquartile range (Q3-Q1) from Q1 or Q3. Data in A, C, E, and F represent mean ± SEM.

We also measured the circadian free calcium rhythms in chloroplasts ([Ca^2+^]_chl_) in Col-0 and *tt4-2* by performing crosses with the *Arabidopsis* MAQ6 line expressing chloroplast-targeted aequorin (Johnson *et al*., 1995; Lenzoni & Knight, 2018). Interestingly, the luminescence of this reporter indicated that [Ca^2+^]_chl_ was elevated in *tt4-2* relative to Col-0, although there was no difference in the amplitude of rhythmic [Ca^2+^]_chl_ (Fig 6C-E). The *tt4-2* seedlings also exhibited a significant reduction in the well-established spike in [Ca^2+^]_chl_ that occurs at the transition from light to dark (Fig. 6F) (Lenzoni & Knight, 2018; Martí Ruiz *et al*., 2018; Pivato *et al*, 2023). Together, these data indicate that flavonoid deficiency in *Arabidopsis* alters Ca^2+^ homeostasis in the chloroplast but not the cytosol. This suggests that a loss of antioxidant potential could affect crosstalk between H_2_O_2_ and Ca^2+^ signaling pathways (e.g., Bhattacharyya *et al*, 2025) specifically in chloroplasts, a major site of ROS production. This finding also points to retrograde signaling between the chloroplast and the nucleus as a potential target of flavonoid action that indirectly affects *TOC1:LUC* amplitude.

## Discussion

We previously reported that the lack of CHS, the first enzyme in the flavonoid pathway, leads to the dysregulation of core clock gene expression in *Arabidopsis* and increased the amplitude of *TOC1:LUC* (Hildreth *et al*., 2022). These findings indicate that the circadian clock is influenced by flavonoid metabolism, although the underlying mechanism is unknown. In this study, we focused on the change in the amplitude of *TOC1:LUC* to examine this question, hypothesizing that flavonoids, rather than the enzyme itself, are the important mediators of this regulation, and testing which characteristics of flavonoids underlie this effect. We found that antioxidant activity, but not activation of the NPR1 receptor, one of the few identified players in redox modulation of the core clock (Zhou *et al*., 2015), or altered auxin transport were important for the action of flavonoids on *TOC1:LUC* amplitude (Figs. 3-5, EV4). In addition, although changes in cytoplasmic [Ca^2+^] were not detected in plants devoid of flavonoids, the *tt4-2* line did exhibit changes in Ca^2+^ levels in the chloroplast (Fig. 6). Interestingly, our analyses further indicate that the hydroxylation pattern of the B ring is a significant factor in regulating redox homeostasis and clock amplitude (Fig. 4), as for many other physiological processes in plants and in animals.

Many prior reports have highlighted the potent antioxidant potential of dihydroxy B-ring flavonoids as a prime biochemical property of this class of compounds. *In vitro* experiments have consistently shown that quercetin, a dihydroxy B-ring flavonol, has a higher antioxidant activity than kaempferol, its monohydroxylated counterpart (Agati *et al*., 2012; Csepregi & Hideg, 2018; Nakabayashi *et al*., 2014; Pietta, 2000). *In vivo* studies have also noted that dihydroxy B-ring flavonoids have a significant impact on redox homeostasis (Agati *et al*., 2020; Daryanavard *et al*., 2023; Gayomba & Muday, 2020; Hernández *et al*., 2009; Nakabayashi *et al*., 2014; Xu *et al*., 2017). In our study all of the effects of flavonoids on *TOC1:LUC* amplitude appear to rely on the ability of seedlings to produce dihydroxy B-ring flavonoids, minimizing the possibility that other metabolites are involved, such as ubiquinone, for which kaempferol serves as a precursor (Berger *et al*, 2022), or terpenes and glucosinolates, the products pathways that are interconnected with flavonoid metabolism (e.g., Naik *et al*, 2023; Sugimoto *et al*, 2021). It is worth noting that primary metabolites with strong antioxidant activity have also been implicated in control of ROS signaling into the clock (Kotchoni *et al*, 2009; Philippou *et al*, 2020; Román *et al*., 2021) but that little is yet known regarding the mechanism of action. While the exact mechanism behind the modulation of circadian amplitude by flavonoids also remains unclear, our data point to the antioxidant role of dihydroxy B-ring flavonoids, potentially in chloroplast, as the key to this relationship.

Signaling via [Ca^2+^]_cyt_ has also been implicated in the control of clock function in response to light and diverse stresses (Hotta *et al*, 2008; Kidokoro *et al*, 2022; Martí Ruiz *et al*., 2018). While we did not observe changes in [Ca^2+^]_cyt_ in *tt4-2*, we did detect elevated [Ca^2+^]_chl_ in these seedlings. Similar specific changes in [Ca^2+^]_chl_ have been described in *Arabidopsis* seedlings in response to heat and high light stress (Kuang *et al*, 2024; Lenzoni & Knight, 2018). We also observed a reduction in the spike in [Ca^2+^]_chl_ that occurs at the light-dark transition (Fig. 6E), a well-established phenomenon whose functional significance remains unknown (Martí Ruiz *et al*., 2018; Pivato *et al*., 2023). Interestingly, there is an emerging understanding of mechanisms connecting Ca^2+^ and H_2_O_2_ signaling, not only in the cytoplasm (e.g., Bhattacharyya *et al*., 2025), but increasingly in chloroplasts. For example, elevated [Ca^2+^]_chl_ in response to treatment with oxidative stressors, including H_2_O_2_, has been reported in *Arabidopsis* cell cultures, in *Chlamydomonas*, and in the diatom, *Phaeodactylum tricornutum* (Flori *et al*, 2024; Pivato *et al*., 2023; Sello *et al*, 2016). This hints that the elevated H_2_O_2_ in *tt4* could be the driver of altered Ca^2+^ dynamics in the chloroplast. Flavonoids have been found at low levels within both the cytoplasm and chloroplasts (Agati *et al*., 2012; Winkel, 2019), raising the possibility that flavonoids could influence [H_2_O_2_]_chl_ either directly or indirectly, thereby altering [Ca^2+^]_chl_, which could then act through retrograde signaling to affect clock gene expression in the nucleus.

A key to understanding the mechanisms by which flavonoids regulate the amplitude of *TOC1:LUC in vivo* lies with identifying the target proteins. Our data showed that most flavonoid glycosides exhibit little or no rhythmic accumulation in *Arabidopsis* seedlings, despite the strong circadian rhythmicity of flavonoid enzyme expression (Harmer *et al*., 2000; Liebelt *et al*., 2019; Nagel *et al*., 2015). This may be due to differences in flavonoid production across cell types or subcellular localization that are not detected in whole seedling or leaf extracts, or reflect damping of the levels of individual forms due turnover after reacting with ROS. Addressing this possibility will be another important goal of future efforts to characterize the functions of specific flavonoids in influencing circadian rhythmicity in plants.

## Methods

### Plant Materials and Growth Conditions

*Arabidopsis* lines used in this study include the flavonoid mutants *tt4-11* (Salk_020583; Bowerman *et al*., 2012), a backcrossed version of *tt4-2* (Bennett *et al*., 2006), and *tt7-5* (Salk_053394; Bowerman *et al*., 2012) in the Col-0 ecotype, while *npr1-3 TOC1:LUC* (Cao *et al*, 1997; Zhou *et al*., 2016) and aequorin reporter lines MAQ2/MAQ6.8 (Knight *et al*., 1991; Lenzoni & Knight, 2018) were in the Col ecotype. The *tt4-11* and *tt7-5* lines expressing *TOC1:LUC* were generated previously (Hildreth *et al*., 2022).

Seeds were surface sterilized as described previously (Kubasek *et al*, 1992). For all experiments, seeds were sown on 0.8% agar containing 1x MS (Caisson Laboratories, Smithfield, Utah) with 2% sucrose, adjusted to pH 5.7 with 0.1N KOH. Plates were sealed with Nesco film (Karlan Research Products Corp., Cottonwood, AZ) and stratified for 3-4 days at 4°C. Seedlings were then grown in 12h light:12h dark at 22°C under approximately 150 µE LED lights in an E-30B growth chamber (Percival, Perry, IA). For experiments, plates within each desired genotype were selected at random for treatment groups. Experiments were not blinded.

For exogenous flavonoid and other chemical treatments, seeds were sown on agar plates containing 0.1% DMSO (control) or varying concentrations of naringenin, kaempferol, cyanidin (all from Sigma, St. Louis, MO), quercetin (MP Biomedicals, Solon, OH), or NPA (Neta Scientific, Hainesport, NJ) in 0.1% DMSO. KI treatment was performed by sowing seeds on agar plates containing water (control) or 100 µM KI (Sigma, St. Louis, MO). For DPI treatment, seedlings were treated topically with 100 µl of 0, 1, 10, or 30 µM DPI (Sigma, St. Louis, MO) in 0.1% DMSO at dusk on day 6 just prior to initiating luminescence recording.

### Luciferase Reporter Assay

Ten *TOC1:LUC* seeds were sown in a cluster onto 35 mm agar plates and grown in LD for 6 days as described above. For naringenin, DPI, and NPA treatment experiments, seeds were sown in clusters of fifteen. At dusk on day 5, seedlings were treated with 1 mM D-luciferin (potassium salt; GoldBio, St. Louis, MO). Luminescence was recorded for 7 d starting with dusk on day 6 in constant darkness using a LumiCycle 32 luminometer (Actimetrics, Wilmette, IN). Raw data from days 1-7 were baseline detrended and processed using Fast Fourier Transform Non-Linear Least Squares (FFT NLLS) analysis in BioDare2 (Zieliński *et al*, 2014) to determine amplitude and period. Rhythms that could not fit to a period between 18-30 hours were excluded from amplitude or period quantifications.

### Intracellular Ca^2+^ measurements

*tt4-2* was crossed with MAQ2 or MAQ6.8 aequorin reporter lines, visually screened for the flavonoid-deficient phenotype at the F2 stage, and screened for the presence of the aequorin reporter at the F3 stage via polymerase chain reaction (PCR) (primers used for PCR listed in Table EV1) and luminescence measurements. F3 and F4 seeds homozygous for both the flavonoid mutation and the aequorin reporter gene were used for subsequent experiments. [Ca^2+^]_cyt_ and [Ca^2+^]_chl_ rhythms were measured using seedlings expressing cytosolic (MAQ2) or chloroplast-targeted (MAQ6) aequorin reporter genes, respectively. For this, twenty seeds were sown onto 35mm agar plates and grown as described above. At dusk on day 5, seedlings were treated with 10 µM coelenterazine (Invitrogen, Carlsbad, CA). Luminescence recordings began at dusk on day 6 for 7 d. We quantified amplitude and period of the rhythms as described above. Due to high noise in the [Ca^2+^]_chl_ rhythms, the rhythms were first smoothed in Lumicycle Analysis software (Actimetrics, Wilmette, IN) using smooth 30 prior to amplitude quantification.

### Hydrogen Peroxide Assays

For hydrogen peroxide measurements, 25-30 seeds were sown onto 35 mm agar plates and seedlings were grown in LD for 6 days as described above. Plates were kept under constant light or wrapped with aluminum foil for constant darkness starting on day 6 for 24 h before harvesting whole seedlings every 4 h for 24 h. For light/dark experiments, seedlings were harvested every 4 h for 24 h starting on day 7. For flavonoid/chemical treatments, seedlings were transferred to constant darkness on day 6 and collected after 24 h before harvesting seedlings every 4 h for 24 h. Collections in darkness were done under green LED light (∼20 µE). Seedlings were weighed and snap-frozen in 1.5 ml microfuge tubes containing stainless steel beads (2.3 mm diameter, Small Parts, Logansport, IA) and kept frozen at −80°C. Frozen samples were ground to a powder using a TissueLyser II (Qiagen, Germantown, MD) for three 10-sec pulses, submerging tubes in liquid nitrogen between pulses to prevent thawing. H_2_O_2_ levels were measured using the Amplex^TM^ Red assay (Invitrogen) as described elsewhere (Chakraborty *et al*, 2016; Le *et al*, 2015) with minor adjustments. Briefly, for each sample 200 µl of 50 mM sodium phosphate pH 7.4 buffer was added per 30 mg of tissue. Samples were vortexed and then rocked at 4°C for 15 min before centrifugation at 13,000 rpm, 4°C for 5 min. The supernatant was moved to fresh 1.5 ml microfuge tubes and centrifuged again to remove excess debris. The resulting supernatant was diluted 1/5 with 50 mM sodium phosphate pH 7.4 buffer and 25 µl of diluted sample was added to 25 µl of Amplex^TM^ Red working solution in a half area black 96-well plate (Corning, Kennebunk, ME). The reaction was incubated at room temperature for 30 min, and fluorescence (545 nm excitation/590 nm emission) was measured with a BioTek Cytation5 plate reader (Agilent, Santa Clara, CA). H_2_O_2_ content in pmol per mg fresh weight was calculated from absorbance values using a standard curve as described in the assay protocol provided by the manufacturer.

### LC-MS

Samples were generated by sowing 25 sterilized seeds on 60 mm plates containing 1X MS, 2% sucrose overlaid with 30 µm nylon mesh (ELKO Filtering Co.) to prevent interference from agar contamination (Hildreth *et al*., 2020). Plates were incubated at 22°C with a 12h/12h LD cycle for 6 d. On day 7, collections began at dawn (zeitgeber time 0, indicating the onset of light) and continued every 4 h for 24 h. Seedlings were collected in 1.5 ml microfuge tubes containing two stainless steel beads (2.3 mm diameter, Small Parts, Logansport, IA), weighed, snap frozen in liquid nitrogen, and stored at −80°C. Frozen samples were ground to a powder using a TissueLyser II as described above. The powdered samples were extracted in 200 µl methanol with 0.1% formic acid, vortexed, and sonicated for 5 min. The samples were then centrifuged at 13,000 rpm, 4°C for 10 min. and 180 µl of the supernatant was transferred to a fresh 1.5 ml tube. Pelleted material was then re-extracted with 200 µl methanol and the above process was repeated a second time. After pooling both 180 µl aliquots, the samples were dried and stored at −80°C.

Analyses were performed on a Shimadzu LCMS 9030 QToF mass spectrometer interfaced with a LC-40B X3 UPLC, a SIL-40C X3 autosampler (10°C) and a CTO-40C column oven (40°C). A BEH C18 column (2.1 mm × 50 mm, 1.7-μm particle size; Waters) was used for chromatographic separation with solvent A (0.1% formic acid in water) and solvent B (0.1% formic acid in MeOH) at a flow rate of 0.4 ml min^−1^. Solvent conditions began at 2% B and held for 1 min, then a linear gradient to 30% B at 5 min and finally to 98% B at 10 min, which was held for 3 min. The gradient returned to starting conditions with a 0.5 min gradient to 2% B, followed by a 2.5 min hold. Sample injection volumes were 3 µl. Data was collected in both positive and negative modes in separate injections with MS scanning only. A master mix of all samples was prepared and analyzed by DDA and/or MS/MS. The resulting fragmentation patterns were matched to flavonoid glycosides in ReSpeck and MassBank databases using MS-DIAL software (Horai *et al*, 2010; Sawada *et al*, 2012; Tsugawa *et al*, 2015). Data processing was performed using MetaboAnalyst to obtain peak areas for metabolomic features (Pang *et al*, 2022). Raw LC-MS data and flavonoid MS/MS fragmentation patterns are listed in File S2.

### Statistical Analyses

Statistical tests requiring multiple comparisons were performed using one-way ANOVA with Tukey’s Honestly Significant Difference (HSD) post-hoc test using the agricolae package in R with a significant cutoff of p<0.05. Groups found to statistically differ are assigned different letters in order of highest mean, where a>b>c. Groups assigned ab are not statistically different from groups assigned a or b, groups assigned bc are not statistically different from groups assigned b or c, and groups assigned abc are not statistically different from groups assigned a, b, or c. Specific p-values from HSD tests are listed in File S1. Linear regression analysis was performed using the lm function in R. Correlation coefficient of the regression was reported as the adjusted R^2^. To quantify rhythmicity of metabolites, LC-MS values were normalized to those at ZT0 and rhythmicity parameters were quantified using MetaCycle with LS, ARS, and JTK algorithms (Wu *et al*, 2016). Rhythmic metabolites were defined as meta2d_p<0.05.

## Acknowledgments

This work was supported by grants from the National Science Foundation (grant IOB-0820674 to BSJW and GRFP Award DGE-2235205 to ESL) and a Lay Nam Chang Dean’s Discovery Grant from the College of Science at Virginia Tech (to BSJW and SK). Seeds for the *tt4-2* backcrossed line, *TOC1:LUC*, *npr1-3 TOC1:LUC*, and pMAQ2/pMAQ6.8 were generous gifts from Gloria Muday (Wake Forest University), Robert McClung (Dartmouth College), Xinnian Dong (Duke University), and Marc Knight (Durham University), respectively. LC-MS experiments were performed at the Virginia Tech Mass Spectrometry Incubator (VT-MSI).

## Expanded View Tables/Figures

**Table EV1.**
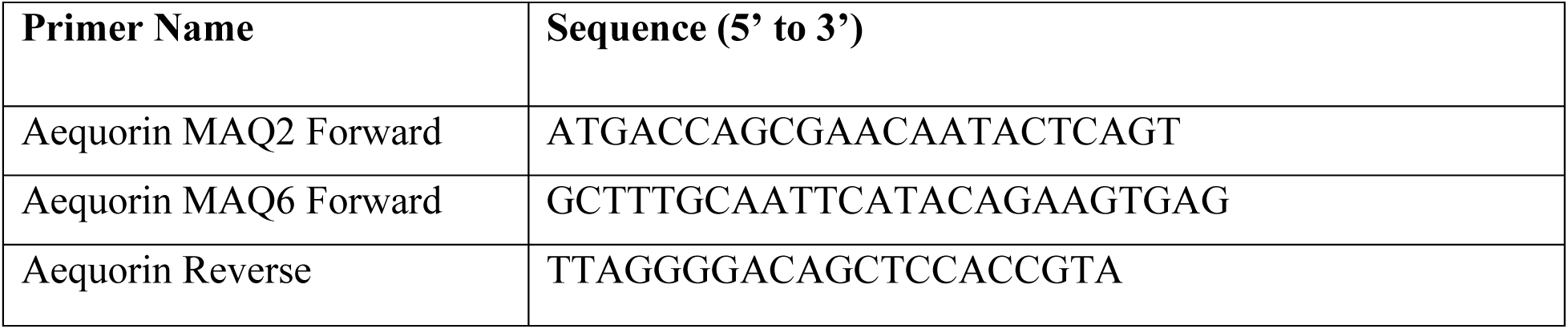
Primers used for genotypic confirmation of Col-0 and *tt4-2* aequorin reporter lines.

**Figure EV1:**
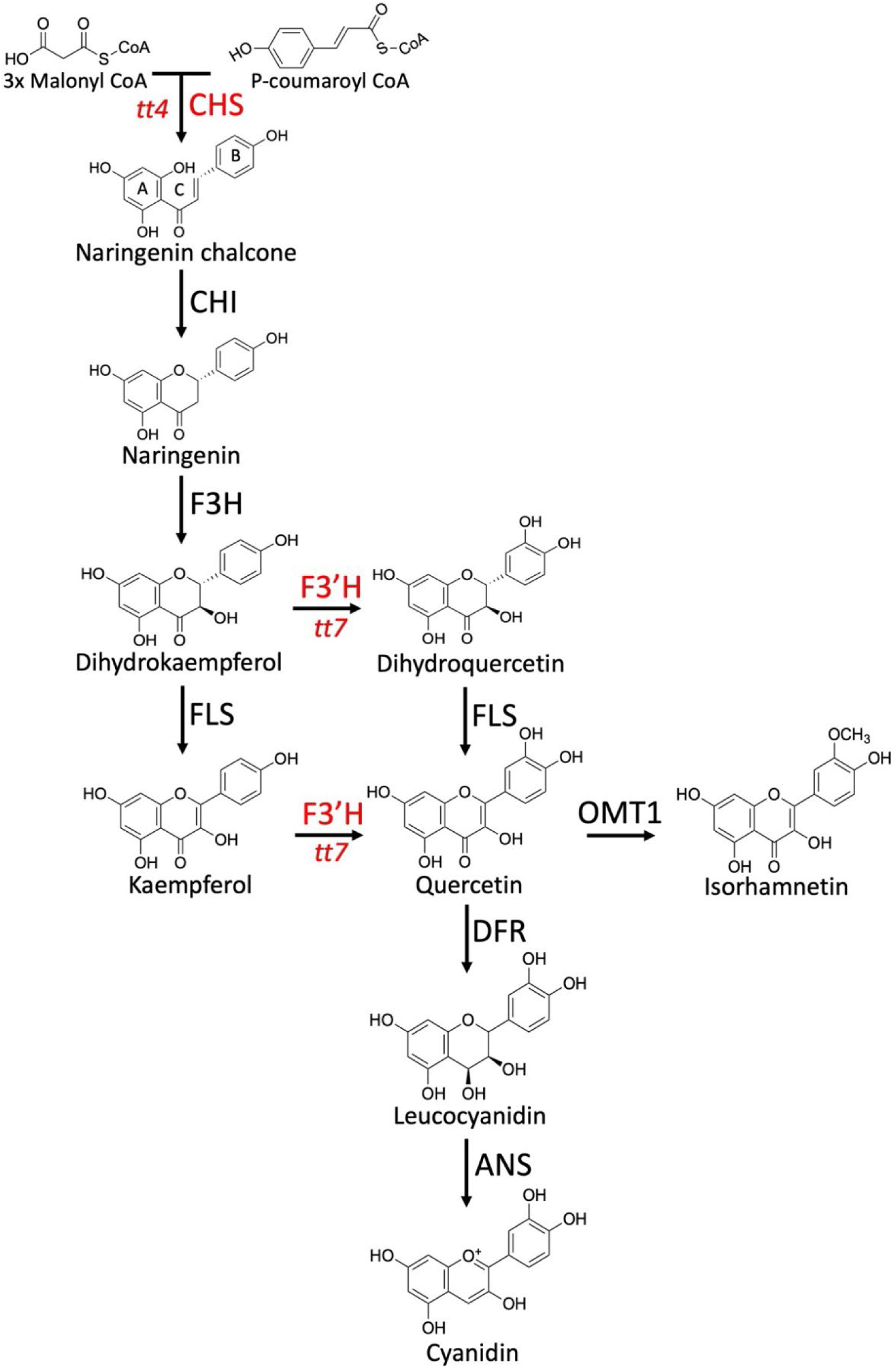
Diagram of flavonoid biosynthetic pathway in *Arabidopsis.* A, B, and C in naringenin chalcone illustrate the convention for naming of the three rings. CHS and F3’H-null lines (*tt4* and *tt7*, respectively) used in this study are shown in red.

**Figure EV2:**
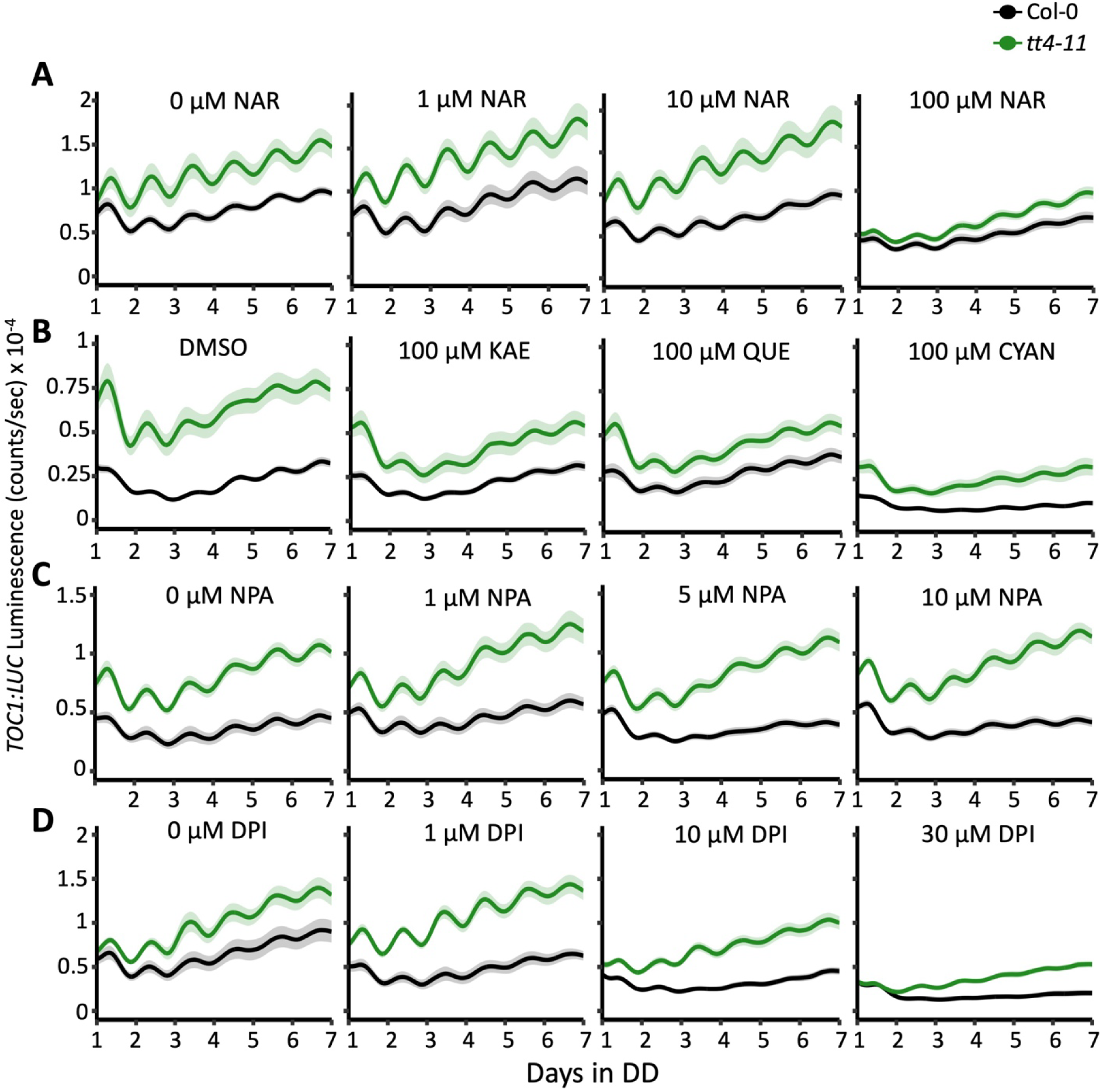
Bioluminescence output of *TOC1:LUC* in Col-0 (black) or *tt4-11* (green) seedlings. (A) Recordings from seedlings grown on 0, 1, 10, or 100 µM naringenin (NAR) (n=8 from two independent experiments), (B) 0.1% DMSO and 100 µM of kaempferol (KAE), quercetin (QUE), or cyanidin (CYAN) (n=12 from three independent experiments), (C) 0, 1, 5, or 10 µM NPA (n=11 from three independent experiments), or (D) 0, 1, 10, or 30 µM DPI (n=6-8 from two independent experiments). Data represents mean ± SEM.

**Figure EV3:**
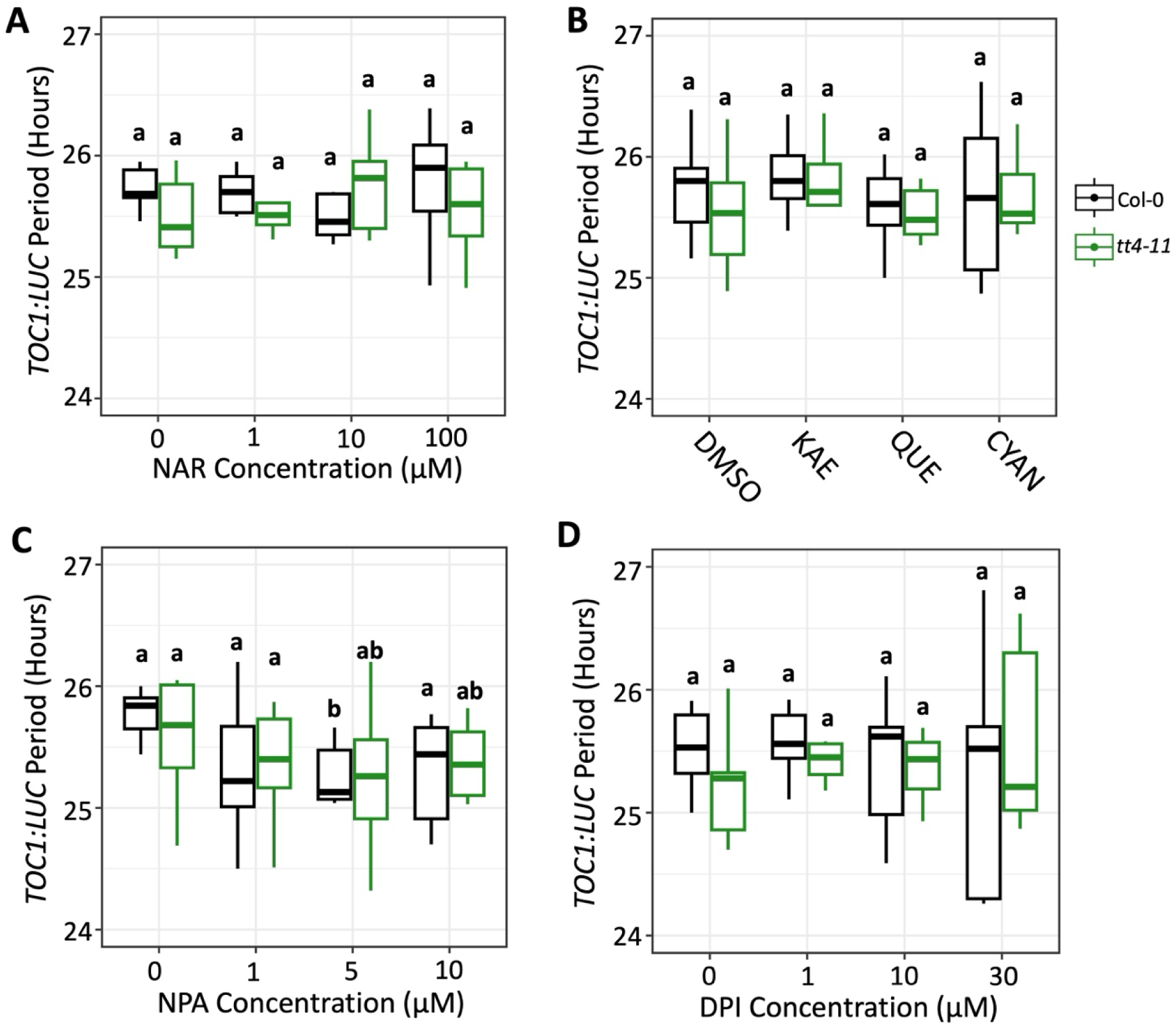
Quantification of *TOC1:LUC* period across chemical treatments. Period of *TOC1:LUC* rhythms in Col-0 (black) or *tt4-11* (green) in the presence of (A) naringenin (NAR) (n= 7-8), (B) kaempferol (KAE), quercetin (QUE), and cyanidin (CYAN) (n= 11-12), (C) NPA (n= 11), or (D) DPI (n= 6-8). Letters represent grouping from one-way ANOVA followed by Tukey post-hoc test with significance cutoff of p<0.05. Boxplot midlines represent median value (Q2), lower and upper line represent 25^th^ (Q1) and 75^th^ percentiles (Q3), respectively. Whiskers represent range of data within 1.5 interquartile range (Q3-Q1) from Q1 or Q3.

**Figure EV4:**
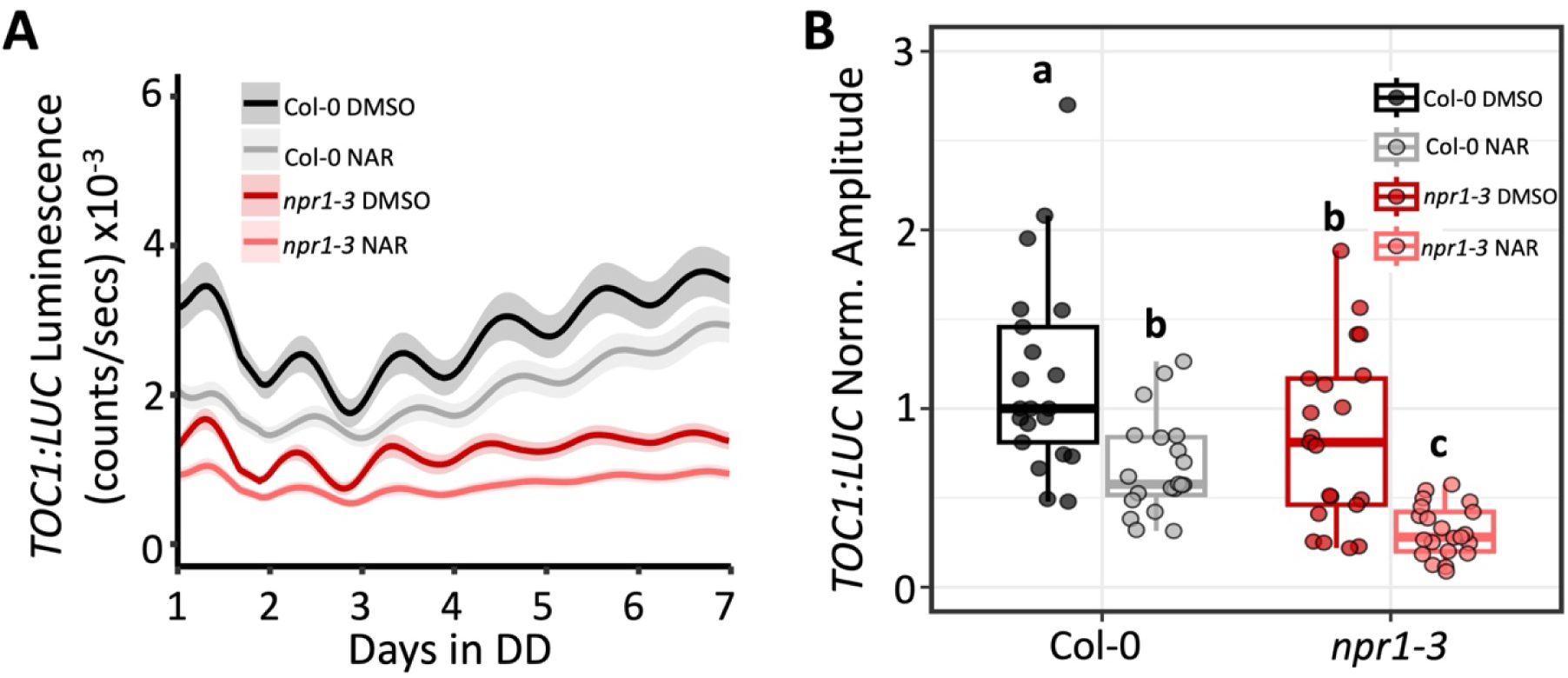
Flavonoids do not regulate circadian amplitude in an NPR1 dependent manner. (A) Bioluminescence output of *TOC1:LUC* in Col-0 (black) or *npr1-3* (red) seedlings grown on 0.1% DMSO or 100 µM NAR (n=20-21 from three independent experiments). (B) Amplitude of *TOC1:LUC* rhythms in Col-0 or *npr1-3* normalized to Col-0 DMSO, which was set to 1. Letters in B represent grouping from one-way ANOVA followed by Tukey post-hoc test with significance cutoff of p<0.05. In B boxplot midlines represent median value (Q2), lower and upper line represent 25^th^ (Q1) and 75^th^ percentiles (Q3), respectively. Whiskers represent range of data within 1.5 interquartile range (Q3-Q1) from Q1 or Q3. Solid line and shading in A, C, D, and E represent mean ± SEM.

